# ESCRT and autophagy cooperate to repair ESX-1-dependent damage to the *Mycobacterium*-containing vacuole

**DOI:** 10.1101/334755

**Authors:** Ana T. López-Jiménez, Elena Cardenal-Muñoz, Florence Leuba, Lilli Gerstenmaier, Monica Hagedorn, Jason S. King, Thierry Soldati

## Abstract

Phagocytes capture invader microbes within the bactericidal phagosome. Some pathogens subvert killing by damaging and escaping from this compartment. To prevent and fight bacterial escape, cells contain and repair the membrane damage, or finally eliminate the cytosolic escapees. All eukaryotic cells engage highly conserved mechanisms to ensure integrity of membranes in a multitude of physiological and pathological situations, including the Endosomal Sorting Complex Required for Transport (ESCRT) and autophagy machineries. In *Dictyostelium discoideum*, recruitment of the ESCRT-III protein Snf7/Chmp4/Vps32 and the ATPase Vps4 to sites of membrane repair relies on the ESCRT-I component Tsg101 and occurs in absence of Ca^2+^. The ESX-1 dependent membrane perforations produced by the pathogen *Mycobacterium marinum* separately engage both ESCRT and autophagy. In absence of Tsg101, *M. marinum* escapes earlier to the cytosol, where it is restricted by xenophagy. We propose that ESCRT has an evolutionary conserved function in containing intracellular pathogens in intact compartments.

## Introduction

After phagocytic uptake, the closely related pathogenic bacteria *Mycobacterium tuberculosis* and *M. marinum* reside in an altered and maturation-arrested phagosome, thereby avoiding its toxic chemical environment^1^, but remaining protected from the cell-autonomous cytosolic defences^2^. This *Mycobacterium*-containing vacuole (MCV) becomes permissive for the bacilli to survive and replicate^3, 4^. However, bacteria access to nutrients is limited. To circumvent this restriction, tubercular mycobacteria damage the MCV and escape to the cytosol. The site of MCV rupture becomes a complex battlefield where various machineries cooperate to repair membrane damage and control cytosolic bacteria. Here, we used the *Dictyostelium discoideum-M. marinum* system to study the role of Endosomal Sorting Complex Required for Transport (ESCRT) and autophagy in membrane repair during both sterile and pathogen-induced damage. We show that the function of ESCRT-III in membrane repair is evolutionarily conserved, that it contributes to the integrity of the MCV and plays an unrecognised role in cell-autonomous defence. We also provide evidence that the ESCRT-III and autophagy pathways act in parallel to repair endomembrane compartments, but differ in their ability to restrict mycobacteria growth in the cytosol of infected cells.

To access the cytosol, mycobacteria make use of a crucial pathogenicity locus, the Region of Difference 1 (RD1), which encodes the ESX-1 system responsible for the secretion of the membranolytic peptide ESAT-6^5^. The membrane perforations produced by ESAT-6 cause MCV rupture and bacterial escape to the cytosol^3, 6, 7^, a step that precedes egress of the bacteria and their dissemination to neighboring cells [reviewed in^8, 9^]. At the site of MCV rupture cells need to discriminate self from non-self as well as from topologically misplaced self molecules. Damage exposes either pathogen-associated molecular patterns (PAMPs) or damage-associated molecular patterns (DAMPs, from the vacuole) to cytosolic machineries that sense them, resulting in the deposition of “repair-me” and “eat-me” signals. Among the latter, the best studied during infection by various intracellular bacteria, including mycobacteria, is ubiquitin. It is conjugated to mycobacterial or host proteins by the E3 ligases NEDD4^10^, Smurf1^11^ and TRIM16^12^ leading to the recruitment of the autophagy machinery to restrict their growth (xenophagy). In addition, mammalian galectins such as Gal-3^12, 13^, Gal-8^14^ and Gal-9^15, 16^ bind to exposed lumenal glycosylations of damaged endosomes^17, 18^ or to components of the cell wall of bacterial pathogens, and thus play a defence role against mycobacterial infection. Although the signaling that leads from these eat-me tags to the recruitment of autophagy is well understood^2, 19^, how the recruitment of the repair machinery(ies) is mediated remains mysterious^20^.

Strikingly, autophagy has been shown to participate in restricting the growth of some bacteria^20, 21^, whilst also contributing to the establishment and maintenance of the compartments containing *Salmonella* Typhimurium^22^ and *M. marinum*^*^20^*^. Recent studies have highlighted the potential of another cytosolic machinery, the ESCRT, in both repair of membranes damaged by sterile insults^23, 24^ and in the control of mycobacteria infection^25^. ESCRT is an evolutionary-conserved machinery, composed of four protein complexes (ESCRT-0, -I, -II, -III), the AAA-ATPase Vps4 and multiple accessory proteins, involved in various processes of membrane remodeling. This is achieved by the ESCRT-III component Chmp4/Snf7/Vps32, which polymerizes in spirals that drive membrane deformation^26^, whereas Vps4 triggers membrane scission and ESCRT-III disassembly^27, 28, 29^. The most studied functions are in the invagination of intralumenal vesicles of multivesicular bodies, which is initiated by the ubiquitination of the cargo to be sorted^30^, in the release of viruses by budding off the plasma membrane^31^ and in the constriction of the cytokinetic bridge during mitosis^32^. Additional roles for ESCRT have been uncovered, such as exosome and microvesicle biogenesis, neuron pruning, removal of defective nuclear pores, and micro- and macroautophagy^33^. Importantly, ESCRT-III has also recently been proposed to mediate repair at a number of membranes such as plasma membrane wounds of less than 100 nm, possibly by surface extrusion^23^, as well as small disruptions in endolysosomes^34^. The ESCRT complexes are implicated in infection of *Drosophila* S2 cells with *M. fortuitum*^35^ and *Listeria monocytogenes*^36^. Follow-up studies showed that depletion of the ESCRT proteins Vps28 or Tsg101 lead to an increase of *M. smegmatis* proliferation in RAW macrophages. These bacteria were found to be highly ubiquitinated at 3 hours post-infection (hpi) in S2 cells, and electron microscopy (EM) inspection revealed that most of them were inside a vacuolar compartment. In the case of *L. monocytogenes*, a mutant unable to secrete the pore-forming toxin LLO, but still expressing two membranolytic phospholipases C, was found to escape more to the cytosol in S2 cells devoid of several ESCRT proteins (Bro/ALIX/AlxA, Vps4 and Snf7/Chmp4/Vps32)^37^. In this context, ESCRT was involved in membrane trafficking and the establishment of the pathogen-containing compartment. However, whether ESCRT plays a role in the repair of damage inflicted by the bacteria remains to be addressed.

Recent reports brought evidence for a role of ESCRT-III in repairing sterile damages to various membranes, such as the plasma membrane (laser wounding, detergents, pore-forming toxins)^23, 24^, the nuclear membrane (laser wounding and confined cell migration)^38, 39^ and endolysosomes (lysosomotropic compounds and silica crystals)^34^. ESCRT-III recruitment to damage at the plasma membrane and lysosomes was proposed to depend on the recognition of a local increase of Ca^2+^ by ALIX and/or ALG2^23, 24, 34^. In all other cases, ESCRT-III is recruited to the disrupted membranes, but the cues and signaling pathways involved remain unclear.

The ESCRT and autophagy machineries are highly conserved in the social amoeba *D. discoideum*^41^. In the last decade, *D. discoideum* has emerged as a powerful model system to study host-pathogen interactions, including with the human pathogens *Legionella pneumophila, Pseudomonas aeruginosa, Vibrio cholerae*, as well as various mycobacteria species such as *M. tuberculosis* and *M. marinum* [reviewed in^40^].

## Results

### The ultrastructure of the MCV at the site of rupture reveals a complex battlefield between *M. marinum* and its host

Previous work demonstrated that the MCV suffers continuous ESX-1-dependent insults and injuries during infection, from the first hour post-entry, when no macroscopic sign of breakage is observed, until about 24 hpi, when the bacteria escape to the cytosol^3^. Ubiquitination is among the first readout of damage and was shown to trigger the recruitment of the classical autophagy machinery components such as Atg18, p62 and Atg8^20^. In order to gain a deeper morphological and ultrastructural insight into the sites of MCV membrane damage during *M. marinum* infection, cells at 24 hpi were subjected to Focus Ion Beam Scanning Electron Microscopy (FIB-SEM) (Fig. 1). This revealed a complex interface between the MCV and the cytosol at the site of bacteria escape. Fig. 1A, B and C show views and 3D reconstructions of bacteria escaping from an MCV and captured by an autophagosome. The zone surrounding the portion of the bacteria in contact with the cytosol shows both discontinuities and highly electron-dense material (Fig. 1C-D). Careful inspection of the samples, together with their 3D reconstruction, revealed that this compact mass was apparently not separated from the MCV content or the host cytosol by any membrane (Fig. 1D-E and Supplementary Movie 1). These observations suggest that bacterial and host factors accumulate at the place of MCV rupture. In macrophages and *D. discoideum*, damaged MCVs and escaping *M. marinum* accumulate ubiquitin^20, 42, 43^. Therefore, we speculate that the dark electron-dense material observed at the areas of MCV rupture might correspond to proteins belonging to the autophagy pathway and possibly other cytosolic machineries, such as the ESCRT, recently implicated in endolysosomal membrane damage repair^34^.

**Figure 1.**
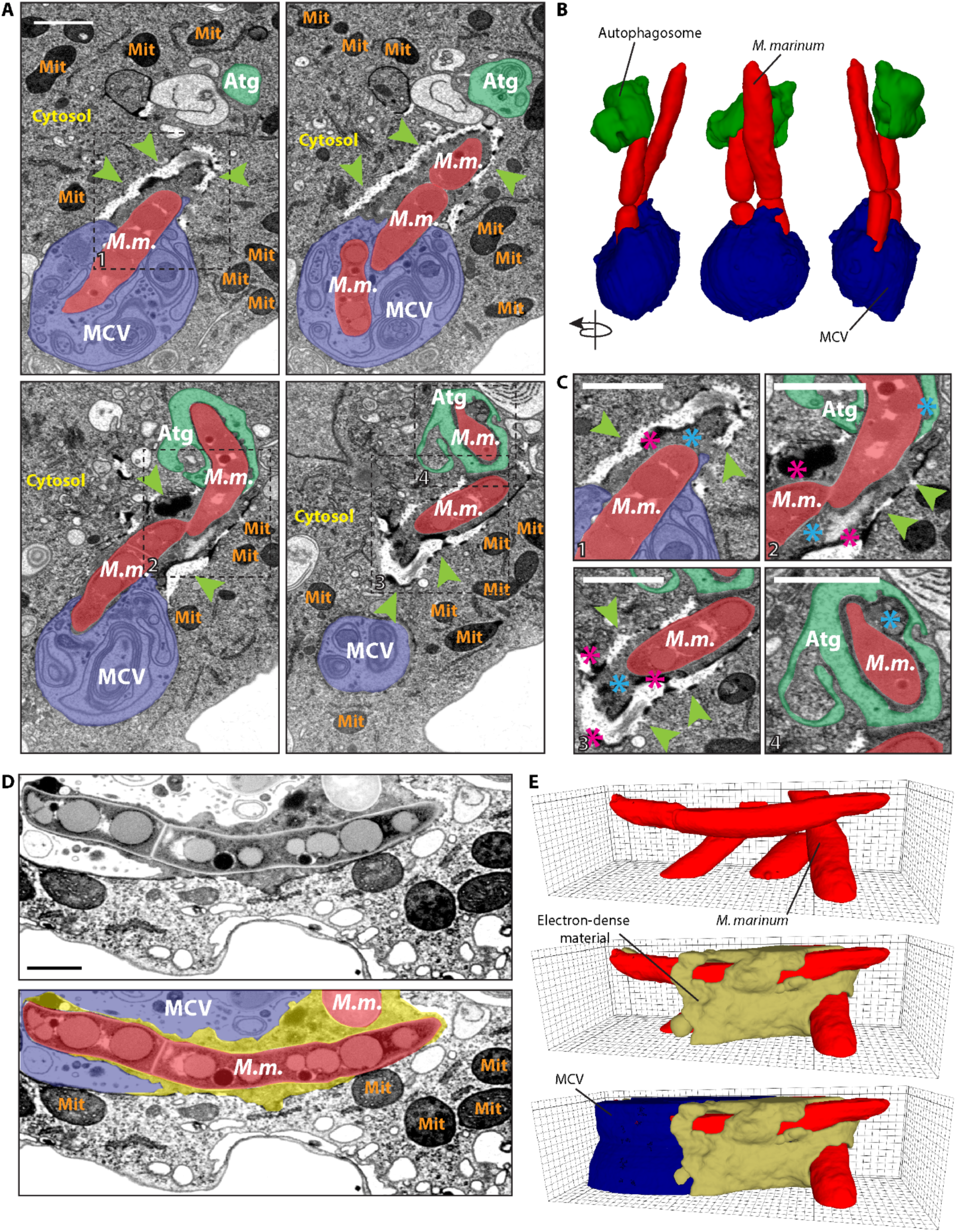
The ultrastructure of the MCV at the point of rupture reveals a complex battlefield between the *M. marinum* and its host. *D. discoideum* was infected with *M. marinum* and fixed at 24 hpi for visualization by FIB-SEM. (A) Sections of the same infected cell, showing a disrupted MCV (blue), *M. marinum* (red) accessing the cytosol and a potential autophagosome (green) engulfing the pole of a cytosolic bacterium. Green arrowheads point to a complex structure with discontinuities and electron-dense boundaries surrounding the cytosolic *M marinum*. Squares delimitate the areas of interest magnified in C. (B) 3D reconstruction of the FIB-SEM stack shown in A. (C) The cytosolic bacteria were surrounded by a structure (green arrowheads) with very electron-dense boundaries (pink asterisks). The cytosolic material between the bacteria and this structure or the autophagosome was slightly more electron-dense than the rest of the cytosol (blue asterisks) (D). Section of a cell, showing a disrupted MCV (blue), *M. marinum* (red) and the dark electron-dense material surrounding the sites of escape (blue asterisks, yellow). (E) 3D reconstruction of the FIB-SEM stack shown in D (see also Movie S1). Abbreviations: *M. m*., *M. marinum*; MCV, *Mycobacterium*-containing vacuole; Mit, Mitochondria; Atg, Autophagosome. Scale bars, 1 µm.

### ESCRT is recruited to the MCV upon *M. marinum*-induced damage

One of the first host responses to membrane damage is the ubiquitination of the bacilli and the broken MCV, followed by recruitment of the autophagy machinery to delay escape to the cytosol^20^. To test whether the ESCRT machinery is also recruited to damaged MCVs, cells expressing the ESCRT-I component GFP-Tsg101, the ESCRT-III effector GFP-Vps32 or the ATPase Vps4-GFP were infected with wild-type (wt) *M. marinum* or *M. marinum* ΔRD1 (Fig. 2A-D). All three proteins were recruited to MCVs containing wt *M. marinum*, but were significantly less so in cells infected with the attenuated *M. marinum* ΔRD1, which causes very limited membrane damage and escapes very inefficiently to the cytosol. The ESCRT-positive structures comprised small foci, patches of several micrometers and even rings that were observed to slide along the length of the bacteria compartment (Fig. 2E and Supplementary Movie 2). These structures seemed to become larger at later time-points (Fig. 2A, 24 and 31 hpi), suggesting increased recruitment upon cumulative damage as infection progresses. Careful 3D inspection at late time-points revealed that GFP-Vps32 patches always surrounded the MCV, but were not in its lumen (Fig. 2F). During membrane remodeling in mammalian cells, the ESCRT-III complex can be recruited to biological membranes via several pathways, one of the most studied relies on the ESCRT-I component Tsg101^44^. Importantly, in cells lacking Tsg101, GFP-Vps32 structures were significantly reduced at early times of infection (Fig. 2G-H), which may indicate that *M. marinum*-induced damage triggers one or more pathways of ESCRT-III recruitment to the MCV. Altogether, we conclude that the ESCRT-III is recruited to the MCV in an ESX-1 dependent manner, consistent with a role in membrane repair.

**Figure 2.**
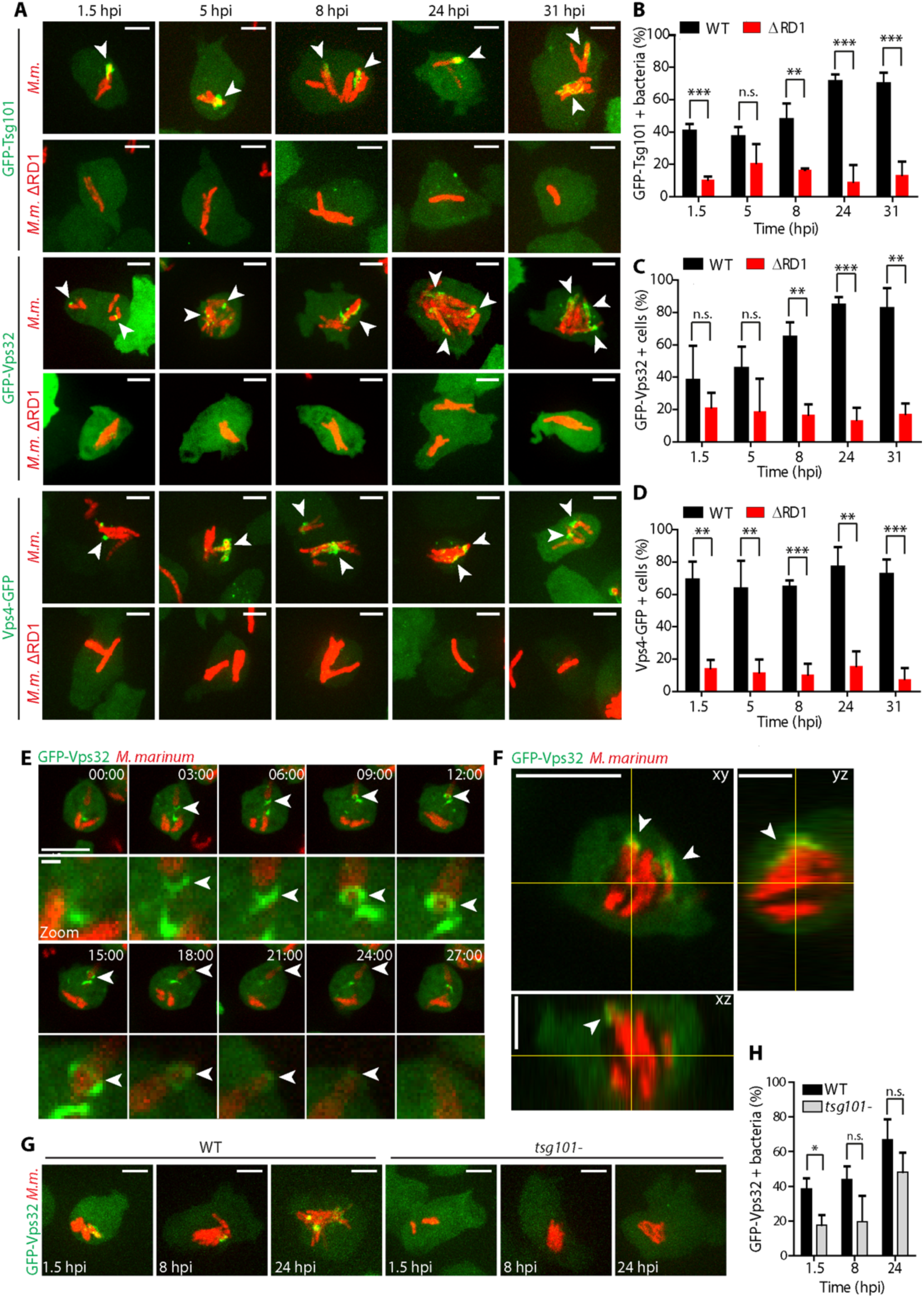
Recruitment of ESCRT components to the MCV upon *M. marinum* induced damage. (A) *D. discoideum* expressing GFP-Tsg101, GFP-Vps32 or GFP-Vps4 were infected with *M. marinum* wt or *M. marinum* ΔRD1, and z-stacks were acquired at 1.5, 5, 8, 24 and 31 hpi. Maximum projections are shown. GFP-Tsg101, GFPVps32 and Vps4-GFP structures (white arrows) appeared in the vicinity of the MCV containing *M. marinum* wt (red), and to a lesser extent around *M. marinum* ΔRD1 (red). Scale bars, 5 µm. (B-D) Quantification of GFPTsg101 structures (patches and foci) and GFP-Vps32 and Vps4-GFP structures (patches and rings) in the vicinity of *M. marinum* wt or *M. marinum* ΔRD1. Plots show the mean and standard deviation (GFP-Tsg101 1.5, 8, 24,31 hpi N=3, 61≤n≤246; 5 hpi N=2, 56≤n≤167, GFP-Vps32 N=3 32≤n≤145; Vps4-GFP N=3, 64≤n≤233). (E) *D. discoideum* expressing GFP-Vps32 were infected with *M. marinum* wt (red) and monitored by time-lapse microscopy every 3 min (see also Movie S2). Maximum projections of the same cell are shown (time indicated in the top right corner). GFP-Vps32 rings formed and appeared to move along the bacterium (arrows). Bottom panels show insets focused on the ring structures (arrows). Scale bars, 10 µm and 1 µm for the insets. (E) Section of a z-stack showing the recruitment of GFP-Vps32 to the vicinity of the MCV (bacteria in red) at 31 hpi. Projections of the xz and yz planes are shown. Scale bar, 10 µm. (G) *D. discoideum* wt or *tsg101-* expressing GFP-Vps32 were infected with *M. marinum* (in red) and z-stacks were acquired at 1.5, 8 and 24 hpi. Maximum projections are shown. GFP-Vps32 was recruited to a lesser extent to the MCV in *tsg101-* cells. Scale bars, 5 µm. (H) Quantification of GFP-Vps32 structures formed in the vicinity of *M. marinum* in wt or *tsg101-* cells. The plot shows the mean and standard deviation (N=3, 76≤n≤154). Two-tailed *t*-tests were performed.

### ESCRT-III and autophagy are recruited to the disrupted MCV at spatially distinct sites

In order to gain a deeper insight on the GFP-Vps32 structures observed during *M. marinum* infection, the precise localization of GFP-Vps32 on the MCV was analyzed. The MCV membrane was visualised by the presence of the ammonium transporter AmtA-mCherry, or antibodies against the predicted copper transporter p80^20^ (Fig. 3A-B). MCVs with a continuous staining for p80 or AmtA-mCherry (Fig. 3A-B, 1.5 hpi) were not associated with GFP-Vps32. On the contrary, compartments that displayed disrupted staining for p80 or AmtA-mCherry presented numerous GFP-Vps32 patches at the sites of membrane wounds (Fig. 3A-B, 8, 24 and 31 hpi). Close inspection of these images revealed that, at sites of GFP-Vps32 recruitment, the damaged MCV membrane was sometimes invaginated towards the lumen of the compartment, away from the cytosol (Fig. 3B, 31 hpi). Time-lapse microscopy enabled tracking of the GFP-Vps32 structures associated with the MCV, indicating direct assembly onto the membrane of the compartment rather than delivery via pre-existing structures (Fig. 3C). GFP-Vps32 structures remained associated with the MCV for several minutes (Fig. 3C and Supplementary Movie 3).

**Figure 3.**
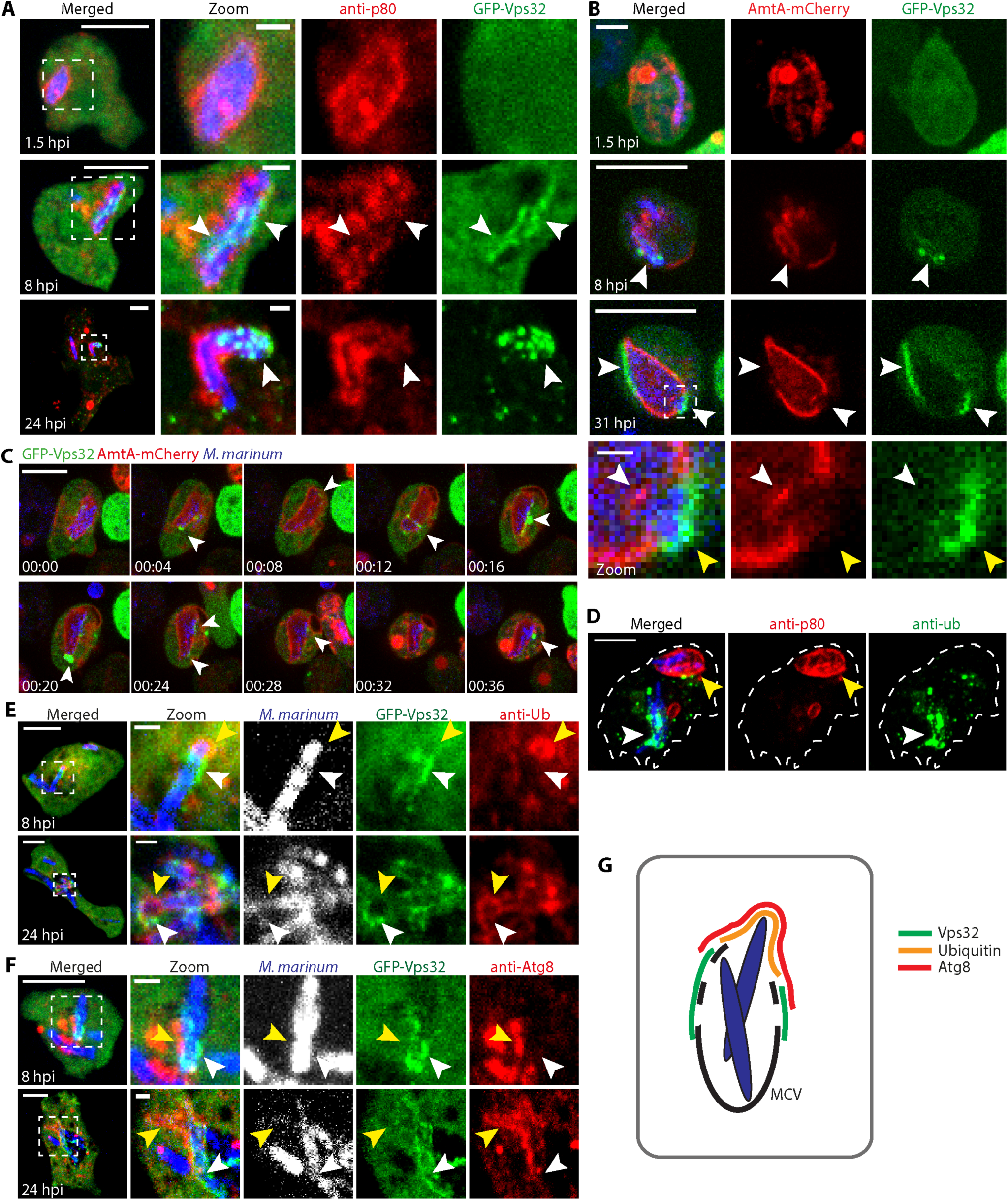
GFP-Vps32 and GFP-Atg8 are recruited to the disrupted MCV at spatially distinct sites. (A) *D. discoideum* expressing GFP-Vps32 were infected with *M. marinum* (blue) and fixed for immunostaining against p80 to label the MCV (red). At 1.5 hpi, no GFP-Vps32 structure was visible at intact MCVs. At 8 and 24 hpi, GFP-Vps32 foci and patches appeared in the vicinity of the bacteria, where the MCV was disrupted (arrows). Scale bar, 10 µm and 1 µm for the insets. (B-C) *D. discoideum* expressing GFP-Vps32 and AmtA-mCherry were infected with *M. marinum* (blue) and monitored by time-lapse microscopy. (B) Still images show GFP-Vps32 patches on the disrupted MCV (arrows). Insets highlight the topology of GFP-Vps32 recruitment (yellow arrow) to the damaged MCV, where the membrane appeared to invaginate towards the lumen of the MCV (white arrow). (C) Time-lapse showing the dynamics of association of GFP-Vps32 to the AmtA-mCherry labelled MCV (see also Movie S3). Scale bars, 10 µm and 1 µm for the insets. (D) *D. discoideum* was infected with *M. marinum* and fixed for immunostaining at 24 hpi. Only bacteria (blue) that have escaped the MCV (p80, red) showed ubiquitin structures (green). Scale bar, 5 µm. (E-F) *D. discoideum* expressing GFP-Vps32 were infected with *M. marinum* (blue) and fixed for immunostaining at 8 and 24 hpi to visualize ubiquitin or Atg8 (red). GFP-Vps32 and ubiquitin or Atg8 were recruited to the same macroscopic region of the MCV, but they did not perfectly colocalise. GFP-Vps32 formed patches devoid of ubiquitin or Atg8 staining (white arrows). Vice versa, ubiquitin and Atg8 appeared in areas where no GFP-Vps32 was observed (yellow arrows). Scale bar, 5 µm and 1µm for the insets. (G) Schematic representation of the Vps32, ubiquitin and Atg8 recruitments at the damaged MCV.

*M. marinum* in intact MCVs stained by p80 are rarely ubiquitinated, contrary to bacteria in the cytosol (Fig. 3D). Therefore, we wondered whether Vps32 would be recruited at sites of ubiquitination, together with the main autophagy marker Atg8. Remarkably, all three proteins localized at disrupted MCVs, but the level of colocalisation of Vps32 with ubiquitin (Fig. 3E) or Atg8 (Fig. 3F) was limited. Instead, GFP-Vps32 seemed to be recruited more proximally to the membrane remnants of the MCV than ubiquitin and Atg8, which predominantly decorated the bacteria poles fully exposed to the cytosol. Besides, at the boundary between the zones enriched in GFP-Vps32 and Atg8, GFP-Vps32 was more proximal to the bacteria, possibly indicating an earlier recruitment (Fig. 3E-G). Taken all together, recruitment of ESCRT-III proteins to the *M. marinum* MCV seems to happen earlier and at different places than the autophagic recognition of the bacteria, suggesting that ESCRT-III and the autophagy pathway might play separate functions in repair and additionally in xenophagic capture.

### Differential spatial and temporal recruitment of ESCRT and autophagy upon sterile damage

Mammalian ESCRT and autophagy machineries localize to damaged membranes for the repair of wounds and removal of terminally incapacitated organelles, respectively^23, 24, 34, 38, 39^. To test whether components of both machineries were also involved in membrane repair in *D. discoideum*, cells expressing GFP-Tsg101, GFP-Vps32 or Vps4-GFP, as well as GFP-Atg8 were subjected to membrane damaging agents, such as the detergent digitonin or the lysosome-disrupting agent Leu-Leu-O-Me (LLOMe) (Fig. 4). Digitonin inserts first into the sterol-rich plasma membrane and then, upon endocytosis, reaches the endosomes. Consistent with this, digitonin initially induced at the plasma membrane dots and crescent-shaped structures of GFP-Vps32 and Vps4-GFP (Fig. 4A and Supplementary Movie 4). After a few minutes, dispersed foci appeared throughout the cytoplasm, suggesting progressive disruption of endomembranes. In agreement with a role of ESCRT-III in repair, small foci of the ESCRT-I component Tsg101 were also observed in the vicinity of the plasma membrane with a similar timing, supporting a role upstream of the recruitment of the ESCRT-III effectors. In contrast, treatment with digitonin did not lead to the recruitment of GFP-Atg8 to the plasma membrane, but it remained present in autophagosomes (Fig. 4A and Supplementary Movie 4). These results support a role for ESCRT but not autophagy in plasma membrane repair of this type of wound. On the other hand, LLOMe induced the formation of both ESCRT-(GFP-Tsg101, GFP-Vps32 and Vps4-GFP) and autophagy-(GFP-Atg8) positive structures at the periphery of lysosomes labelled with fluorescent dextran (Fig. 4B and Supplementary Movie 5). The structures were diverse in morphology and dynamics. Whereas at 2.5 min GFP-Tsg101, GFP-Vps32 and Vps4-GFP appeared as discrete foci surrounding lysosomes, GFP-Atg8 formed a more continuous ring that became apparent only 5 min later. This spatial appearance and partial temporal segregation suggest an independent involvement of ESCRT-III and autophagy during lysosome damage. To confirm and extend the involvement of ESCRT-III in membrane repair in *D. discoideum*, cells expressing GFP-Vps32 or Vps4-GFP were monitored while exposed to other sterile damage. Purified recombinant ESAT-6 and the lysosomotrophic agent glycyl-L-phenylalanine 2-naphthylamide (GPN) induced similar structures as digitonin and LLOMe (Supplementary Fig. 1). We noticed that, the structures formed by GFP-Vps32, known to build the polymers that deform membranes^26^ were large and intense (Fig. 4 and Supplementary Fig. 1), and especially long-lived on injured lysosomes, where several foci remained in close apposition to dextran-labelled compartments for several minutes (Fig. 4 and Supplementary Fig. 2). Vps4-GFP structures were less obvious but also long-lived (Fig. 4 and Supplementary Fig. 1). In contrast, GFP-Tsg101 structures were less intense and short-lived (Fig. 4), consistent with its upstream role in recruiting the membrane-remodelling ESCRT-III.

**Figure 4.**
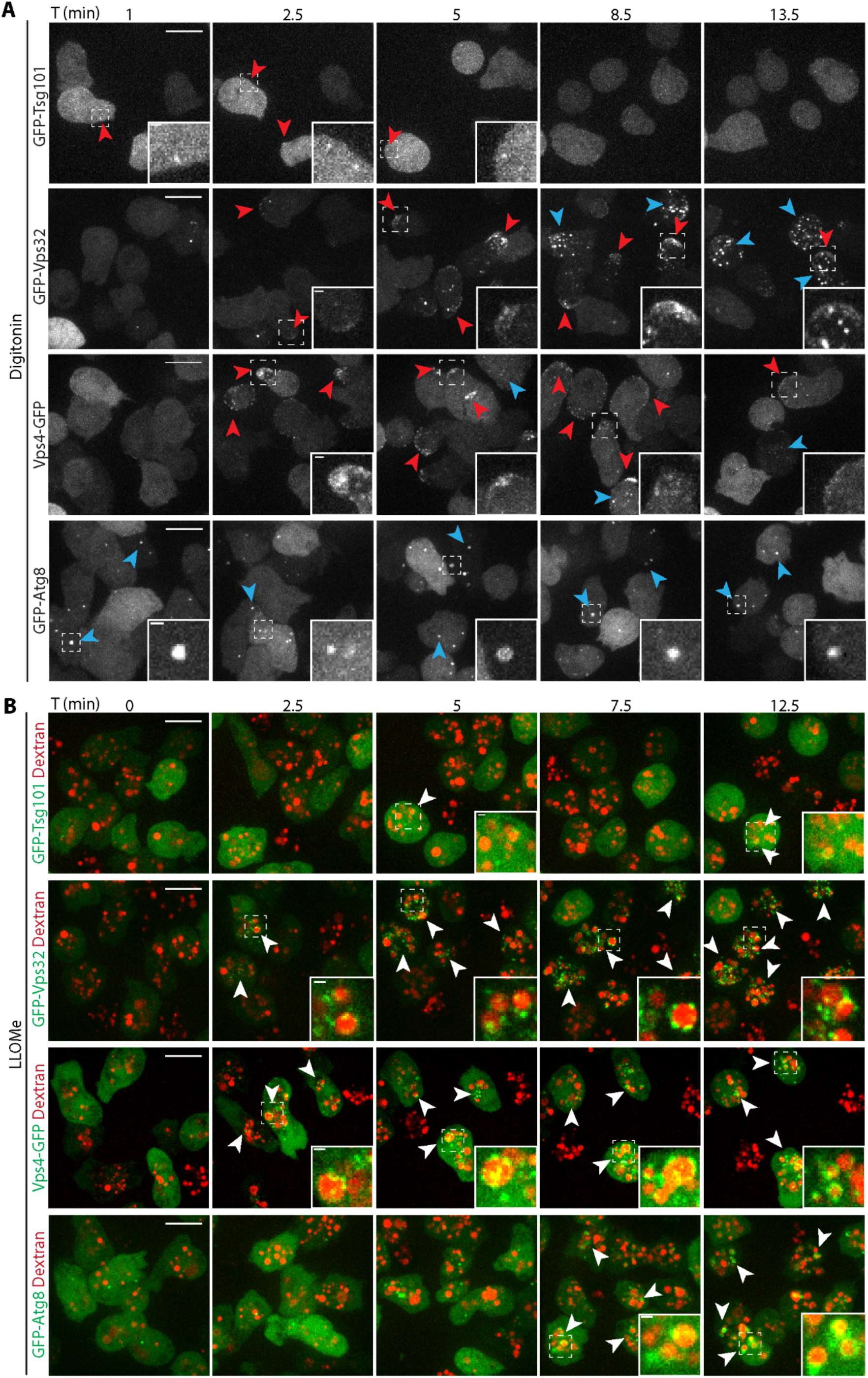
Differential spatial and temporal recruitment of ESCRT components and Atg8 upon sterile damage. *D. discoideum* expressing GFP-Tsg101, GFP-Vps32, Vps4-GFP or GFP-Atg8 were subjected to membrane disrupting agents and monitored by time-lapse imaging. (A) Treatment with digitonin led to the appearance of the three ESCRT-components but not GFP-Atg8 at the plasma membrane within minutes (see also Movie S4). (B) Cells were incubated with 10 kDa fluorescent dextran for at least 3 h to label all endosomal compartments and treated with LLOMe (see also Movie S5). Punctate ESCRT-structures appeared at the periphery of the lysosomes as early as 2.5 min after the addition of LLOMe. More diffuse, ring-like GFP-Atg8 structures were visible about 5 min later. Scale bars, 10 µm or 1 µm for the insets.

### Mechanistic characterization of ESCRT-III recruitment at the sites of sterile damage

To confirm that the ESCRT-III structures formed at the site of membrane repair, cells expressing GFP-Vps32 were treated with digitonin in the presence of fluorescently-labelled Annexin V to reveal exofacially exposed phosphatidyl-serine (PS) (Fig. 5A-B and Supplementary Movie 6). The majority of the GFP-Vps32 crescent structures were also labelled with Annexin V (Fig. 5B). The Annexin V-positive structures were released to the extracellular medium, suggesting that damaged membranes were extruded instead of internalized.

**Figure 5.**
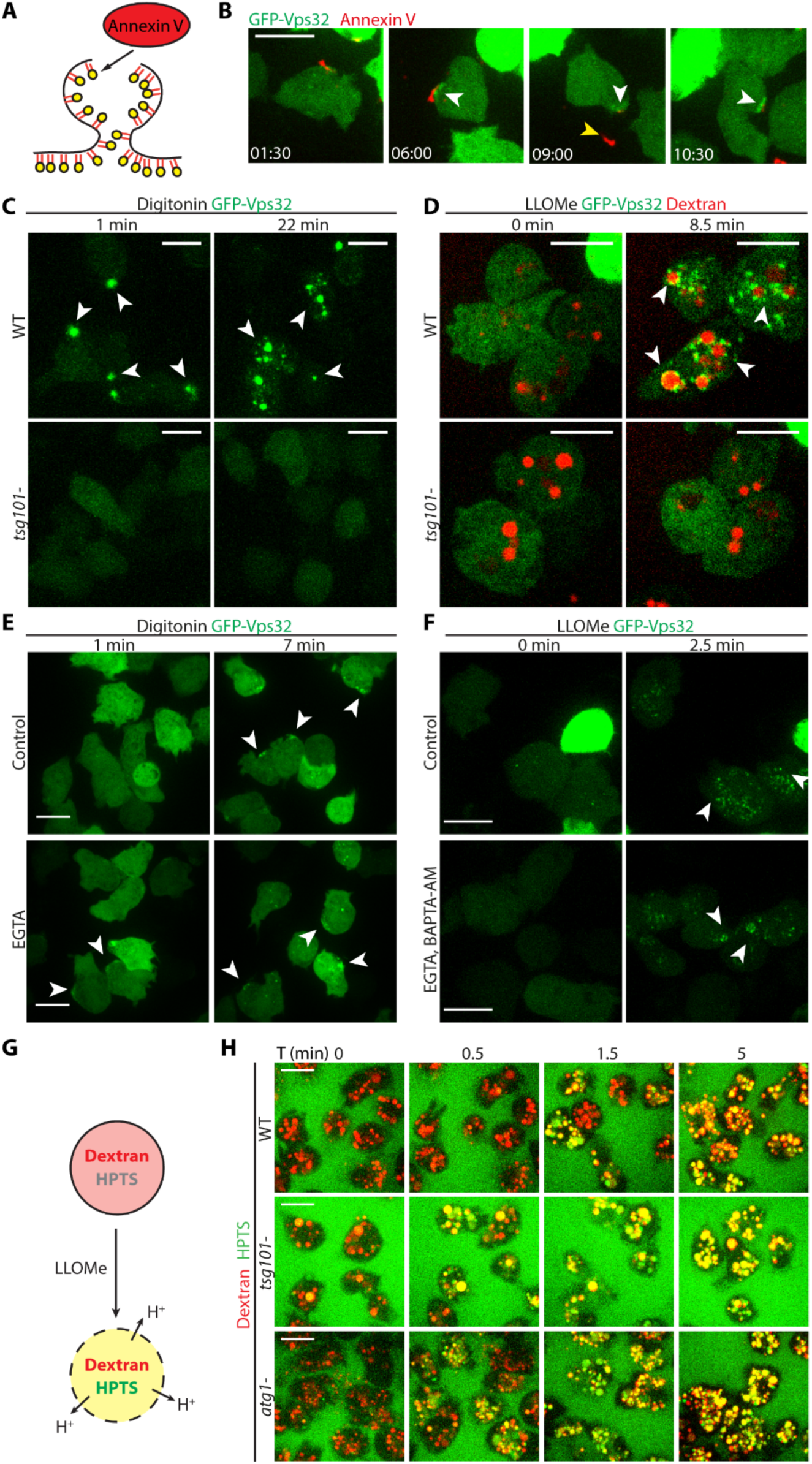
Mechanistic characterization of ESCRT-III and autophagy recruitment and repair of membrane damage. (A) Schematic representation of the experiment shown in B. Annexin V binds to PS exposed upon plasma membrane disruption. (B) *D. discoideum* were incubated with Annexin V Alexa Fluor 594 conjugate in the presence of Ca2+ and then treated with digitonin and monitored by time-lapse imaging (see also Movie S5). GFP-Vps32 structures (in green) appeared in close proximity of Annexin V-positive structures (in red, white arrows). At later times, the Annexin V-positive structure was released into the medium (yellow arrows), leaving a GFP-Vps32 “scar” (white arrow). Time is indicated in the bottom left corner. (C-D) In cells lacking Tsg101, neither digitonin nor LLOMe treatment led to the formation of GFP-Vps32 structures. (D) Endosomes (in red) were labelled with 10 kDa fluorescent dextran for at least 3 h. (E) Cells were incubated with EGTA or mock-incubated, treated with digitonin and monitored by time-lapse microscopy. Neither spatial nor temporal differences in GFP-Vps32 recruitment (white arrows) were observed. (F) Cells were incubated with EGTA and BAPTA-AM or mock-incubated, treated with LLOMe and monitored by time-lapse microscopy. Neither spatial nor temporal differences in GFP-Vps32 recruitment (white arrows) were observed. Scale bars correspond to 10 µm. (G) Schematic representation of the experiment shown in H. Cells were treated for at least 3 h with the Alexa Fluor 647 10 kDa dextran (red) to label all endosomes, together with the 0.5 KDa soluble pH indicator HPTS (green). (H) *D. discoideum* wt, *tsg101-* and *atg1-* were subjected to the experimental procedure depicted in G and monitored by time-lapse imaging (see also Movie S6). Before addition of LLOMe, HPTS was quenched in acidic lysosomes, which therefore appeared in red. HPTS dequenching started after 30 sec in *tsg101-* and *atg1-* cells and after 1.5 min in wt cells.

In mammalian cells, ESCRT-III can be recruited to membranes by at least three mechanisms depending on the identity of the membrane and the specific role exerted by ESCRT [reviewed in^44^]. In *D. discoideum* cells lacking Tsg101, GFP-Vps32 structures formed neither upon digitonin nor LLOMe treatments (Fig. 5C-D), providing a strong evidence that Tsg101 lies upstream of ESCRT-III during membrane repair caused by these types of sterile damage.

It has been proposed that the local increase of intracellular Ca^2+^ upon membrane damage recruits ESCRT-III to the plasma and lysosomal membranes in HeLa cells and myoblasts^23, 24, 34^. To test whether the formation of GFP-Vps32 structures also relied on a Ca^2+^-mediated signaling in *D. discoideum*, cells were treated with digitonin in the presence of the non-permeant Ca^2+^ chelator EGTA, or with LLOMe in the presence of EGTA and the cell-permeant BAPTA-AM (Fig. 5E-F). None of these conditions altered the appearance nor the dynamics of the GFP-Vps32 structures at the wound site. In conclusion, in *D. discoideum*, ESCRT-III recruitment to membranes damaged by these sterile agents appears to be Tsg101-dependent but independent of Ca^2+^ signalling.

### Cells lacking Tsg101 or Atg1 are defective at maintaining lysosome integrity upon LLOMe treatment

To further dissect the functional contributions of the ESCRT-III and autophagy machineries to the repair of wounds inflicted by LLOMe, cells were incubated with a mixture of two fluid-phase markers: the 10 kDa pH-insensitive Alexa Fluor 647 dextran and the 0.5 kDa pH sensor 8-Hydroxypyrene-1,3,6-trisulfonic acid, trisodium salt (HPTS), which is quenched at pH < 6.5 (Fig. 5G-H and Movie S7). Around 5 min after LLOMe addition, the HPTS fluorescence increased drastically and synchronously in the lysosomes of wt cells, indicating proton leakage from the compartments. In the autophagy mutant *atg1*-, which are defective in MCV/endomembrane repair^20^, the fluorescence dequenching happened significantly faster, a sign of earlier proton leakage. Interestingly, in *tsg101*-cells, the switch in fluorescence also happened earlier, again suggesting as for the *atg1*-cells a defect in membrane repair in these mutants.

### Deficiencies in membrane repair lead to earlier escape of *M. marinum* from the MCV

To decipher the role of ESCRT during infection, cells lacking Tsg101 or the accessory proteins AlxA and the AlxA interactors Alg2a and Alg2b were infected and examined by EM. In wt cells, *alxA-* or *alg2a-/b-* mutants, the membrane of the MCV in close vicinity to the bacilli escaping the compartment was even and smooth (Supplementary Fig. 3A, E-G). However, in the *tsg101-* mutant, rough and “bubbling” membrane structures were observed (Supplementary Fig. 3B-C), suggesting cumulating membrane damage. In all cases, escaping bacteria were surrounded by the highly electron-dense material already described in Fig. 1.

It was shown that in the *atg1-* mutant, *M. marinum* escapes earlier from the MCV, accumulates ubiquitin but proliferates unrestrictedly in a cytosol devoid of a bactericidal xenophagy pathway^20^. We reasoned that, if the ESCRT-III machinery were involved in repair of the MCV, then, in the *tsg101*-mutant, bacteria might access the cytosol and become ubiquitinated earlier. The percentage of ubiquitinated *M. marinum* at 8 hpi was significantly higher in the *tsg101-* (75.5 ± 4.3%) than in wt cells (40.3 ± 11.6%, Fig. 6A-B and Supplementary Fig. 4C). In agreement with the increased ubiquitination of bacteria, *M. marinum* also colocalized more with Atg8 in the *tsg101*- (Fig. 6C-D and Supplementary Fig. 4D). Although the percentage of ubiquitinated bacteria in *tsg101-* cells was close to that observed in the *atg1*- and *atg1-tsg101*- double mutants (84.6 ± 3.3% and 89.3 ± 7.5%, respectively), the extent of ubiquitin decoration on the bacteria was very different (Fig. 6A-B and Supplementary Fig. 4C). Whereas in cells lacking Tsg101 ubiquitin formed foci or patches around *M. marinum*, in cells devoid of autophagy bacteria were more densely coated with ubiquitin (Fig. 6A and Supplementary Fig. 4C). This accumulation is probably due to the fact that ubiquitinated bacteria cannot be targeted to autophagic degradation in the *atg1-* mutant, but autophagy is still functional in the *tsg101*-mutant.

**Figure 6.**
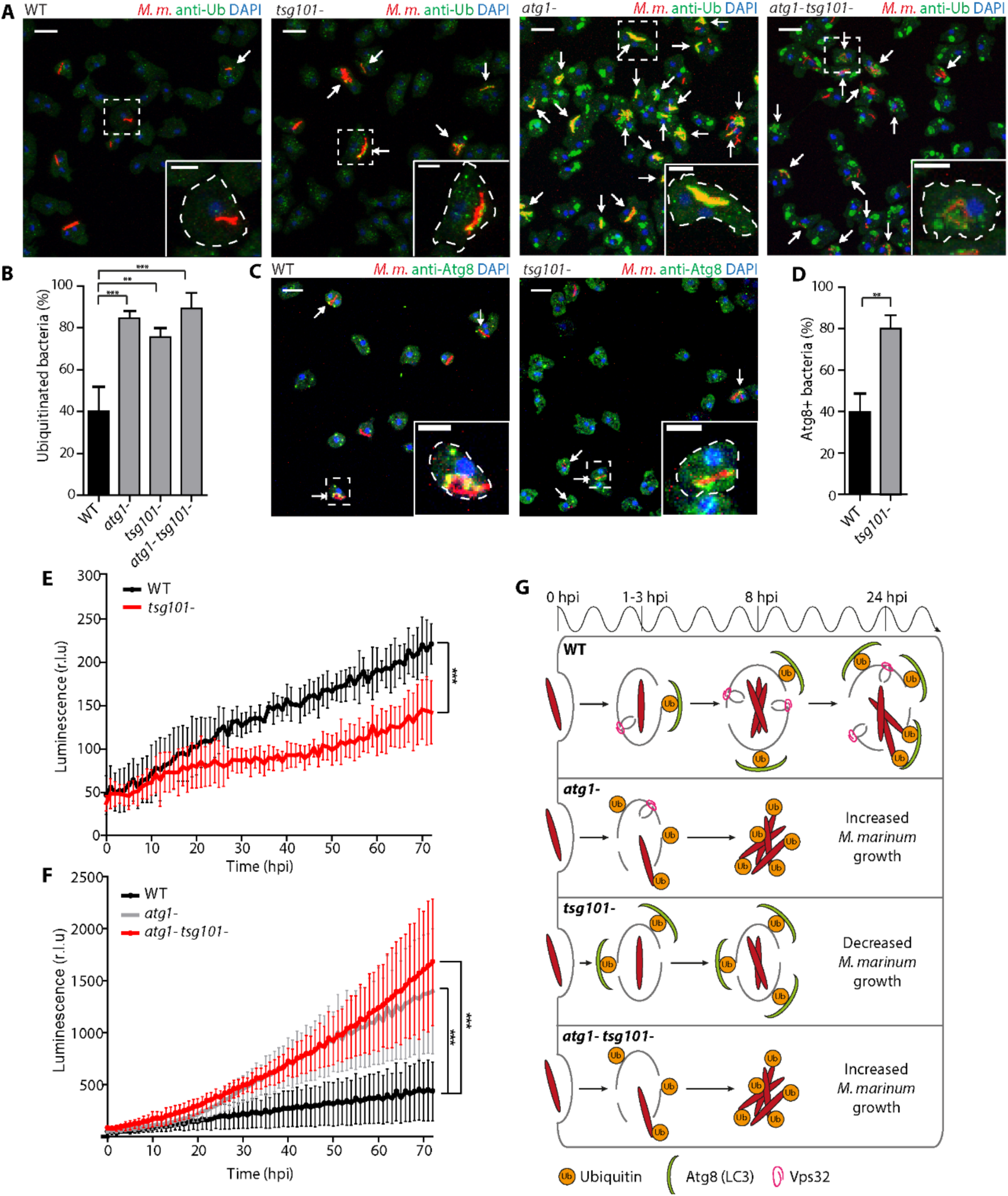
Deficiency in membrane repair leads to earlier escape of *M. marinum* from the MCV and growth restriction by autophagy. (A) *D. discoideum* wt or mutant (*atg1*-, *tsg101*-, *atg1*-*tsg101*-) were infected with *M. marinum* and fixed for immunostaining at 8 hpi (*M. marinum* in red, ubiquitin in green, DAPI in blue). Arrows point to ubiquitinated bacteria. Scale bars, 10 µm and 5 µm for the insets. (B) Quantification of the proportion of ubiquitinated bacteria or bacterial microcolonies. The plot shows the mean and standard deviation (WT N=6, n=306; *atg1-* N=3, n= 251; tsg101-N=3, n= 93; N=3, n= 215). (C) Wt or *tsg101-* cells were infected with *M. marinum* and fixed for immunostaining at 8 hpi (*M. marinum* in red, Atg8 in green, DAPI in blue). Arrows point to bacteria decorated with Atg8. Scale bars, 10 µm and 5 µm for the insets. (D) Quantification of the proportion of bacteria or bacterial microcolonies decorated with Atg8. The plot shows the mean and standard deviation. (WT N=3, n=70; *tsg101-* N=3, n= 63). Two-tailed *t*-tests were performed. (E-F) Wt or mutant *D. discoideum* were infected with luminescent *M. marinum* and intracellular bacterial growth was monitored in a plate reader over 72 hpi. *M. marinum* growth was restricted in the *tsg101-* mutant, whereas it hyperproliferates in the *atg1-* and the double *atg1-tsg101-* mutant. Plots represent the mean and standard deviation of N=3 independent experiments. Two-way ANOVA was performed. (G) Model of autophagy and ESCRT-III involvement in MCV repair and restriction of *M. marinum* intracellular growth. In wt *D. discoideum, M. marinum* induces ESX-1-dependent injuries in the MCV membrane. This leads to the separate recruitment of the ESCRT-III and autophagy machineries to repair the damage, but nevertheless the bacteria are able to access the cytosol at later stages of infection (24 hpi). In the *atg1-* mutant, *M. marinum* access the cytosol earlier, despite the membrane repair exerted by ESCRT-III. Bacteria accumulate ubiquitin and proliferate unrestrictedly in the cytosol devoid of a functional autophagy pathway. In the *tsg101-* mutant, membrane damage not repaired by ESCRT-III leads to an increase of ubiquitination and recruitment of the autophagy machinery, resulting in restriction of *M. marinum* growth. In a double *atg1-tsg101-* mutant, lack of both ESCRT-III and autophagic membrane repairs leads to an earlier access of bacteria to the cytosol, which accumulate ubiquitin and hyperproliferate as in the single *atg1-* mutant.

Given that both ESCRT-III and autophagy are involved in the biogenesis of MVBs and autophagosomes, respectively, which rely at least partially on the recognition of ubiquitinated cargoes, we monitored the morphology of endosomes, as well as the levels of ubiquitination, in non-infected ESCRT and autophagy mutants (Supplementary Fig. 4). In the *atg1-* and *atg1-tsg101-* mutants accumulation of high levels of ubiquitinated material was observed, in agreement with the inability of these mutants to degrade it by autophagy. In *tsg101*-cells, only a minor increase of ubiquitin was observed in endosomal compartments (Supplementary Fig. 4A-B), as already reported^45^, which does not explain the more frequent and larger ubiquitin decorations around *M. marinum* in these cells (Fig. 6A).

In yeast and mammallian cells devoid of some ESCRT proteins, ubiquitinated cargoes are not properly sorted into MVBs and accumulate on the limiting membrane^46^. Therefore, to confirm that the increase in ubiquitination observed during infection of the *tsg101*-mutant was due to MCV damage and bacteria access to the host cytosol, and not to failed endocytic cargo sorting, we monitored the colocalization of bacteria with GFP-tagged perilipin (Plin). Plin is a lipid droplet protein that binds the cell wall of *M. marinum* as soon as the bacteria access the cytosol^47^. Like in *atg1*-cells^47^, recognition of cytosolic *M. marinum* by GFP-Plin was higher in *tsg101*- and *atg1*-*tsg101*-compared to wt cells (Fig. S4E-F), confirming the earlier bacteria escape from the MCV in cells lacking a functional ESCRT machinery. Altogether, these results suggest that both Tsg101 and Atg1 trigger separate membrane repair pathways and restrict *M. marinum* access to the cytosol during infection.

Since we have shown that Tsg101 is not essential for ESCRT-III recruitment to the damaged MCV (Fig. 2G-H), but has an important role in repairing the MCV and constraining bacteria escape (Fig. 6 and Supplementary Fig. 4), we wondered whether the accessory proteins AlxA and Alg2a/b, also known to recruit ESCRT-III, were involved in the repair of the MCV. In cells lacking Alg2a/b, the percentage of ubiquitinated *M. marinum* was comparable to that in its respective parental strain (43.3 ± 15.0% and 50.2 ± 11.4% respectively, Supplementary Fig. 5A-B and E) and, similarly, the degree of Atg8 colocalization with the bacteria remained lower in the *alg2a-/b-* mutant (54.70 ± 9.5%, Supplementary Fig. 5C-D and F). On the contrary, 81.8 ± 9.9% of bacteria were ubiquinated in cells devoid of AlxA, which correlated with a higher but not significant increase of Atg8 recruitment to the bacteria (72.7 ± 13.0%, Supplementary Fig. 5A-F). This suggests that AlxA, together with Tsg101 but not Alg2a/b, contributes to the ESCRT-III-mediated repair of the MCV.

### Impairment of ESCRT or autophagy has a distinct impact on *M. marinum* intracellular growth

To study how the ESCRT pathway may impact the outcome of *M. marinum* infection, the ESCRT mutants were infected with luminescent *M. marinum*^48^ and intracellular bacterial growth monitored^49^ (Fig. 6E-F and Supplementary Fig. 5G-H). *M. marinum* luminescence increased around 5-fold in wt *D. discoideum* in the course of 72 h, reflecting sustained intracellular growth. In the *atg1-* mutant, since bacteria escape earlier to a cytosol that is devoid of xenophagic defense, *M. marinum* grew better (Fig. 6F), as already described^20^. *M. marinum* proliferation in both ESCRT mutants *alxA-* and *alg2a-/b-* was similar to that in the wt (Supplementary Fig. 5G-H), suggesting no crucial involvement of these proteins in the infection course.

Importantly, loss of Tsg101 significantly suppressed *M. marinum* growth compared to wt cells (Fig. 6E). Interestingly, in the double mutant *atg1-tsg101-*, bacterial luminescence increased substantially, reaching similar levels as in the single *atg1*-mutant. Therefore, a functional autophagy pathway is necessary to control bacterial burden in the *tsg101*-mutant indicating that without ESCRT-III-mediated MCV repair the bacteria become more accessible to degradation by xenophagy. Taken together with the previous results on ubiquitination and Atg8 recruitment (Fig. 6 and Supplementary Fig. 5), we conclude that Tsg101 and AlxA but not Alg2a/b participate in the ESCRT-III-mediated repair of the MCV damage and thus absence of these proteins enables an earlier escape of *M. marinum* to the cytosol. In the case of *tsg101-*, this leads to the early recruitment of the autophagy machinery, which restricts bacterial growth.

## Discussion

While most intracellular bacterial pathogens reside in a vesicular compartment where they exploit the host resources, a few bacteria have adapted to translocate to the host cytosol. The dynamics of escape to the cytosol varies depending on the pathogen. For instance, while *Shigella* and *Listeria* trigger an early escape, *Salmonella* and mycobacteria program a partial and/or delayed escape^8^. This is the case of *M. marinum*, which disrupts the MCV thanks to the membranolytic activity of ESAT-6, a small bacterial peptide secreted by the ESX-1 system^50^. Perforation of the MCV implies first the leakage of host and bacterial factors contained in its lumen and, eventually, bacteria access to nutrients in the cytosol, which must be sensed and restricted by the host. 3D EM inspection of infected *D. discoideum* cells revealed a very complex interface between *M. marinum* and the host cytosol at the site of MCV rupture (Fig. 1), suggesting a dynamic and complex interplay between bacterial and host factors. Here, we show that the two highly conserved ESCRT-III and autophagy pathways contribute to the repair of the MCV membrane, delaying the escape of *M. marinum* to the cytosol.

Like its mammalian homologs^23, 24, 34, 38, 39^, the *D. discoideum* ESCRT proteins Tsg101, Vps32 and Vps4 localized to injuries both at the plasma membrane and endomembranes upon damage by distinct chemical agents such as digitonin and LLOMe (Fig. 4A-B). Addition to the extracellular medium of ESAT-6 led to the formation of comparable Vps32 and Vps4 foci (Supplementary Fig. 1). Importantly, *M. marinum* infection also leads to the appearance of Tsg101, Vps32 and Vps4 foci, patches or rings in the vicinity of the MCV (Fig. 2). Consistent with a role in membrane repair, these structures were significantly less abundant upon infection with *M. marinum* ΔRD1, an attenuated mutant that lacks the ESX-1 secretion system and thus has reduced capability of inducing damage at the MCV. In agreement with its ability to form packed spiral polymers on membranes, upon both sterile injuries and infection, GFP-Vps32 structures were generally larger and longer-lived than the Vps4-GFP ones. Consistently, large ring-like structures were observed exclusively with GFP-Vps32 (Fig. 2E). Time-lapse microscopy revealed that GFP-Vps32 was recruited from the cytosolic pool, likely polymerized at wounds of the MCV, and remained associated with the MCV for several minutes (Fig. 3C).

The wounds inflicted by membrane disrupting agents such as LLOMe [less than 5 nm^51^] may be of comparable size to the ones caused by the mycobacterial membranolytic peptide ESAT-6 [4.5 nm^6^] and thus lead similarly to the recruitment of the ESCRT-III repair machinery. However, the sustained insults and cumulative damage inflicted by *M. marinum*^*^52^*^ likely results from the constitutive secretion of ESAT-6 and also depends on PDIMs^53, 54^. Together, they probably generate heterogeneous and expanding wounds that are harder to resolve. This may explain why the recruitment of GFP-Vps32 to sterile damage depends strictly on Tsg101 (Fig. 5C-D), whereas during *M. marinum* infection this dependency is partial (Fig. 2G-H), because other ESCRT recruiting pathways likely act simultaneously. In addition, we propose that cumulative damage by *M. marinum* would eventually overwhelm membrane repair by ESCRT-III and result in recruitment of autophagy, as previously suggested for endosomal damage caused by LLOMe^34^. In the case of sterile damage to lysosomes, autophagy may end up degrading the severely injured compartments by lysophagy^34^. Regarding damage to the MCV, we propose that xenophagy generates a compartment that either kills the mycobacteria or creates a restrictive environment that somehow impairs the growth of the pathogen.

Upon digitonin treatment, GFP-Vps32 colocalized with Annexin V-labelled regions of the plasma membrane that were subsequently shed in the medium (Fig. 5-B). This suggests that plasma membrane repair in *D. discoideum* might be achieved by ESCRT-III-dependent budding and scission of the wound, as in mammalian cells^23, 24^. A similar repair mechanism, in this case by budding the injury towards the lumen, has also been described for the nuclear envelope^39^. We propose that the same process may operate at the MCV, which would partially explain the presence of abundant membranous material in the lumen of the MCV and the putative invagination of the MCV membrane remnants, as observed by EM and live microscopy (Fig. 1, 3B and Supplementary Fig. 3A).

*M. marinum* that accesses the host cytosol becomes ubiquitinated^20^ and coated by Plin^47^. We used ubiquitination and recruitment of GFP-Plin as a proxy to monitor bacterial escape from the MCV in cells lacking Tsg101. Similar to the high ubiquitination previously shown to occur in amoebae deficient for autophagy^20^, large ubiquitin patches were observed at the sites of MCV rupture in *tsg101*-cells (Fig. 6A-B). Consistently, the percentage of bacteria coated by Plin was higher in this ESCRT mutant (Supplementary Fig. 4E-F). These results are consistent with the large ubiquitin patches observed on *M. smegmatis* in macrophages^25^ and with the increased cytosolic bacterial spread observed during *L. monocytogenes* infection in S2 cells^37^. In addition to its role in bacteria restriction, autophagy has been proposed to mediate membrane repair of vacuoles containing *Salmonella*^22^ and *Mycobacterium*^*^20^*^. Moreover, ESCRT-III has been shown to participate in endolysosomal membrane repair independently of lysophagy^34^. Here, we suggest that in *D. discoideum* autophagy and ESCRT-III work in parallel to repair the damaged MCV. It has been shown that ubiquitin serves as an “eat-me” signal that targets cytosolic bacteria to autophagic degradation^2^. Consistent with this, the accumulation of ubiquitin around *M. marinum* in *tsg101-* cells correlated with a proportional increase of Atg8 decoration on the bacteria (Fig. 6C-D), and with a decreased bacterial load (Fig. 6E), contrary to what has been described in RAW macrophages infected with the non-pathogenic, vacuolar *M. smegmatis*, in which depletion of Tsg101 led to bacteria hyperproliferation^25^. In addition, only very limited colocalization between GFP-Vps32 and ubiquitin or Atg8 was observed at damaged MCVs (Fig. 3E-F).

The cues and signals that recruit ESCRT-III to damage in *D. discoideum* are still to be identified. The appearance of ESCRT-III components before ubiquitinated material can be detected at the site of lysosome disruptions^34^ speaks for a ubiquitin-independent mechanism, although it cannot be excluded that ubiquitin might participate in a subsequent reinforcement loop to recruit ESCRT-III. In mammalian cells, several reports have suggested that influx of extracellular Ca^2+^ through the plasma membrane or efflux through endolysosomal membranes are essential for the positioning of ESCRT-III to the site of the injury^23, 24, 34^. The local increase of intracellular Ca^2+^ at the wound site might be recognized directly by ALIX^23^, a multifunctional protein involved in cargo protein sorting into intralumenal vesicles [reviewed in^55^], thereby bypassing the need for ESCRT-0, -I and -II, and recruiting ESCRT-III by direct protein-protein interactions. Alternatively, Ca^2+^ has been proposed to be sensed by ALG2, an ALIX-interacting protein with a penta EF-hand domain, which could promote the accumulation of ALIX, ESCRT-III and the Vps4 complex at the damage site^24, 34^. Strikingly, in *D. discoideum* intra- and extracellular Ca^2+^ chelation did not impair GFP-Vps32 relocation to plasma membrane or lysosome injuries (Fig. 5E-F). Consistent with this result, Ca^2+^ seemed also to be dispensable during MCV repair, since knockout of either *alxA* or *alg2a/b* did not impact intracellular *M. marinum* growth (Supplementary Fig. 5G-H). Altogether, our results suggest that the MCV damage caused by the *M. marinum* ESX-1 secretion system is repaired very robustly and in a multilayered response by both the ESCRT-III and the autophagy pathways of *D. discoideum* in a Tsg101-dependent and Ca^2+^-independent manner. The ability of the ESCRT-III to repair membranes injured by various biological and chemical insults strongly suggests that this is a generic mechanism that operates upon infection by other intracellular pathogens that damage membranes such as *Salmonella, Shigella* and *Listeria*. In addition, the high conservation of the ESCRT machinery identifies this pathway as a novel potential therapeutic target to fight against bacterial infection in humans.

## Materials and Methods

### *D. discoideum* strains, culture and plasmids

*D. discoideum* strains and plasmids are summarized in Supplementary Table 1. *D. discoideum* Ax2(Ka) was axenically cultured at 22°C in Hl5c medium (Formedium) supplemented with 100 U mL^-1^ of penicillin and 100 µg mL^-1^ of streptomycin (Invitrogen). *D. discoideum* JH10 was cultured similarly as Ax2(Ka), with the addition of 100 g/mL of Thymidine (Sigma). Cells were transformed by electroporation and selected with the relevant antibiotic. Hygromycin, G418 and blasticidin were used at a concentration of 50, 5 and 5 µg mL^-1^, respectively. *D. discoideum* Ax2(Ka) *tsg101-* and *atg1-tsg101-* were obtained by transformation with the pSP72 KO vector kindly provided by Dr. L. Aubry^56^. JH10 *alxA-* and JH10 *alg2a-/b-* mutants were kindly provided by Dr. L. Aubrey^57, 58^. Ax2(Ka) cells expressing GFP-Atg8 and AmtA-mCherry were described in^20^ and^47^, respectively. Ax2(Ka) cells expressing GFP-Vps32, Vps4-GFP and GFP-Tsg101 were generated by transformation with the vectors described in Supplementary Table 1. Cells expressing GFP-Plin were obtained upon transformation with the pDNeoGFP-Plin construct described in^59^.

### Mycobacteria strains, culture and plasmids

*M. marinum* strains and plasmids are summarized in Supplementary Table 1. *M. marinum* (M strain) wt and ΔRD1 were kindly provided by Dr. L. Ramakhrisnan^20^. Mycobacteria were cultured in 7H9 (Difco) supplemented with glycerol 0.2% (v/v) (Biosciences), Tween-20 0.05% (v/v) (Sigma) and OADC 10% (v/v) (Middlebrock). mCherry-expressing bacteria were obtained by transformation with pCherry10, which was kindly obtained from Dr. L. Ramakrishnan^60^, and cultured in the presence of 50 µg/mL^-1^ hygromycin (Labforce). Luminiscent bacteria harbored the pMV306::*lux* plasmid^48^ and were cultured in presence of 50 µg mL^-1^ kanamycin (AppliChem).

### Infection assay

Infections were performed as previously described^61^. In brief, *M. marinum* bacteria were washed in Hl5c and centrifuged onto adherent *D. discoideum* cells. After additional 20-30 min of incubation, extracellular bacteria were washed off and the infected cells resuspended in Hl5c containing 5 U mL^-1^ of penicillin and 5 µg mL^-1^ of streptomycin (Invitrogen). For live microscopy, mCherry expressing or unlabeled bacteria were used. To monitor bacteria intracellular growth in *D. discoideum*, luciferase-expressing *M. marinum*^20, 49^ were used.

### Intracellular growth measurement

Growth of luminescent bacteria was measured as described previously^20, 49^. Briefly, dilutions of 0.5-2.0 × 10^5^ *D. discoideum* cells infected with *M. marinum* pMV306::*lux* were plated on a non-treated, white F96 MicroWell plate (Nunc) and covered with a gas permeable moisture barrier seal (Bioconcept). Luminescence was measured for 72 h at 1 h intervals with a Synergy Mx Monochromator-Based Multi-Mode Microplate Reader (Biotek). The temperature was kept constant at 25°C.

### Live fluorescence microscopy

Infected or uninfected *D. discoideum* were plated in 35 mm Glass Bottom Dishes (MatTek) or in 4-wells µ-slides (Ibidi). To induce damage in *D. discoideum* membranes, cells were incubated with 2.5-5 µM of digitonin (Sigma), 5 mM of LLOMe (Bachem), 200 µM of GPN or 60 mg mL^-1^ of purified recombinant ESAT-6 (BEI Resources). To visualize the lumen of endocytic compartments, 0.5 mg mL^-1^ of 70 KDa TRITC-Dextran or 10-15 µg mL^-1^ of 10 kDa Alexa Fluor 647 Dextran (Molecular Probes) were added at least 3 h prior to visualization of the sample. To detect neutralization of endocytic vesicles, 0.2 mM of 524 Da HPTS (Molecular Probes) was added at least 3 h prior to the visualization of the sample. To detect plasma membrane damage, 5 µM of Annexin V Alexa Fluor 594 conjugate (Molecular Probles) was added to cells in the presence of 2.5 mM Ca^2+^. To chelate Ca^2+^, 5 mM of EGTA (Fluka) or 5 mM of EGTA and 250 µM of BAPTA-AM (Sigma) were added at least 20 min before imaging. Bacteria not expressing a fluorescent reporter were stained with 12.5 µM of Vybrant DyeCycle Ruby stain (Molecular Probes) directly on the µ-dish prior to microscopic inspection. Live microscopy was performed with a Spinning Disc Confocal Microscope [Intelligent Imaging Innovations Marianas SDC mounted on an inverted microscope (Leica DMIRE2)] with a glycerin 63x or an oil 100x objective. Generally, z-stacks from 3 to 10 slices of 1µm or 1.5 µm were acquired. For time-lapse experiments, images were acquired from every 15 s to several min. Image analysis was performed using ImageJ.

### Cell fixation and immunofluorescence stainings

Samples were fixed in ultracold methanol as already described^62^. Briefly, *D. discoideum* cells on coverslips were quickly fixed by immersion in −85°C methanol using a Freezing Dewar (FH Cryotec, Instrumentenbedarf Kryoelektronenmikroskopie). Subsequent immunofluorescence staining was performed as described^62^. Rabbit anti-Atg8 was previously described^20^, anti-p80^63^ was purchased from the Geneva Antibody Facility (http://www.unige.ch/antibodies), the anti-Ub (FK2) monoclonal antibody was from Enzo Life Sciences^20^. Nuclei were stained with DAPI (Molecular Probes). Cells were embedded using ProlongGold antifade (Molecular Probes). Images were acquired with a LSM700 or LSM780 microscope (Zeiss) using an oil 63x objective. Image analysis was performed using ImageJ.

### Transmission Electron Microscopy (TEM)

Sample preparation for TEM was performed as described in^64^. Briefly, *D. discoideum* cells were fixed in a 6 cm dish in 2% (w/v) glutaraldehyde in Hl5c for 1 h and stained with 2% (w/v) OsO4 in imidazole buffer 0.1 M for 30 min. Cells were detached with a cell scraper and washed 3 times with PBS. Subsequent sample preparation was performed at the Pôle Facultaire de Microscopie Ultrastructurale (University of Geneva). Samples were incubated in 2% (v/v) of Milloning buffer and rinsed with distilled water. Then, they were incubated in 0.25% (w/v) uranyl acetate overnight and rinsed with distilled water. Samples were dehydrated using increasing concentrations of ethanol, then in propylene oxide for 10 min and finally embedded in 50% Epon-propylene oxide for 1h, followed by incubation overnight in pure Epon. Samples were embedded in 2% agar for subsequent sectioning in an ultramicrotome and placed on TEM grids. Finally, sections were visualized in a Tecnai 20 electron microscope (FEI Company, Eindhoven, The Netherlands).

### Focus Ion Beam Scanning Electron Microscopy (FIB-SEM)

Initial sample preparation was performed similarly as for TEM and sent to the Pôle Facultaire de Microscopie Ultrastructurale (University of Geneva). Subsequent contrast enhancement, dehydration and resin embedding was performed as described in^65^ Samples were visualized in a Helios DualBeam NanoLab 660 SEM (Fei Company, Eindhoven, The Netherlands). 3 D reconstitutions were performed using the LimeSeg plugin from ImageJ.

### Statistics

All graphs were plotted and statistical analysis were performed using Prism. In all quantifications, the mean and standard deviation is shown. Two-tailed t-test or 2-way ANOVA was used (n.s: non-significant, *: p-value < 0.05, **: p-value < 0.01, ***: p-value < 0.001).

## Acknowledgements

We would like to acknowledge Dr. Laurence Aubry for her generous gift of antibodies and *D. discoideum* ESCRT mutant strains, Dr. Sonia Arafah for the generation of the *tsg101*- mutant and preliminary work on intracellular growth measurements, Ms Iryna Nikonenko and the PFMU platform of the Centre Medical Universitaire for processing the samples for TEM and FIB-SEM, and the Bioimaging Platform of the Faculty of Sciences. LG and MH were supported by a grant from the Deutsche Forschungsgemeinschaft (HA3474/3-2), and JSK by a Royal Society University Research Fellowship (UF140624). The TS lab is supported by grants from the Swiss National Science Foundation (310030_149390 and 310030_169386) and the SystemsX.ch initiative grant HostPathX.

## Authors contributions

Conceived and designed the experiments: ATLJ, TS. Performed the experiments: ATLJ, FL, ECM. Analysed the data: ATLJ, ECM. Contributed reagents/material/analysis tools: ATL, FL, LG, MH, JSK. Wrote the original draft: ATLJ, TS. Reviewing & editing: ATLJ, ECM, MH, JSK, TS.

